# Ligand Strain Energy in Large Library Docking

**DOI:** 10.1101/2021.04.06.438722

**Authors:** Shuo Gu, Matthew S. Smith, Ying Yang, John J. Irwin, Brian K. Shoichet

**Author notes:** contributed equally (Shuo Gu, Matthew S. Smith).

## Abstract

While small molecule internal strain is crucial to molecular docking, using it in evaluating ligand scores has remained elusive. Here, we investigate a technique that calculates strain using relative torsional populations in the Cambridge Structural Database, enabling fast pre-calculation of these energies. In retrospective studies of large docking screens of the dopamine D4 receptor and of AmpC β-lactamase, where close to 600 docking hits were tested experimentally, including such strain energies improved hit rates by preferentially reducing high-scoring decoy molecules that were strained. In a 40 target subset of the DUD-E benchmark, we found two thresholds that usefully distinguished between ligands and decoys: one based on the total strain energy of the small molecules, and one based on the maximum strain allowed for any given torsion within them. Using these criteria, about 75% of the benchmark targets had improved enrichment after strain filtering. Relying on pre-calculated population distributions, this approach is rapid, taking less than 0.04 second to evaluate a conformation on a standard core, making it pragmatic for pre-calculating strain in even ultra-large libraries. Since it is scoring function agnostic, it may be useful to multiple docking approaches; it is openly available at http://tldr.docking.org

## INTRODUCTION

In large library docking screens, hundreds-of-millions to billions of molecules, each in multiple conformations, are sampled for complementarity to a protein binding site. While low energy conformations predominate, inevitably some high-energy conformations are sampled. In a foundational study, Tirado-Rives and Jorgensen showed that it was often possible to find high energy conformations of small molecules in docking, and that the relative energies of these conformations were difficult to rank, even with fairly high-level quantum mechanics.^1^ The authors concluded that there were inherent errors in docking—here from ligand strain but also from other terms—that were outside the limits of binding affinity ranges docking was likely to sample among its hits (from mM to mid-nM, or about 5 kcal/mol). This made accurate rank ordering in docking unlikely in general, and in particular made the improvement of docking scores by the addition of higher order terms, here ligand strain, problematic.

While docking will remain, for the foreseeable future, a method that cannot reliably rank-order molecules from a large library screen, we wondered if ligand strain could nevertheless improve docking in its primary goal, as a categorizing technique that separates a small fraction of plausible new ligands from a much larger library of molecules unlikely to bind.^2–5^ High-ranking docked molecules can adopt strained conformations, typically owing to high-energy torsion angles. Ideally, such strained conformations should not be sampled, but the need to explore many conformations to achieve favorable fits, combined with necessarily approximate conformational energies and the screening of large, diverse libraries, has often meant that such strained conformations are in fact modeled. Perversely, these strained conformations can often score better in a protein site than a lower energy, unstrained conformation.^6^ This in turn can crowd out unstrained, more favorable molecules, lowering the ability of docking to separate true ligands from false positives. Were such strained conformations removed, it would improve docking hit rates, even if including strain, per se, in the energy score might not measurably improve rank ordering by affinity, as argued by Tirado-Rives and Jorgensen.

Unfortunately, few methods for calculating conformational energies now meet the demands of large-library docking, where hundreds-of-millions to billions of molecules must be assessed,^7–9^ and several hundred thousand must be evaluated even after the initial docking campaign has completed (post-filtering). The current approaches for strain energy assessment can be organized into three groups: quantum mechanical (QM), molecular mechanics (MM), and database-torsional methods. In *ab initio* calculations,^10–13^ torsional energies are calculated using one of several QM basis sets. For example, Rai et al generated torsion scan profiles of neutral fragments using DFT (B3LYP).^14^ Based on the QM calculations at discrete torsion angles, they interpolated the values into a continuous function to estimate the torsion strain energy. This is not yet feasible at scale, and so the method was converted into a semi-knowledge-based approach. Even so, the method still demands *ab initio* calculation of torsional terms for many new molecules, and remains time consuming. In molecular mechanics^15^ approaches, force fields including OPLS, CHARMM, and AMBER^16–19^ are used to calculate small molecule strain energies; this can often demand the calculation of new parameters, especially for large and diverse docking libraries. Molecular mechanics methods are less accurate than QM methods, since they involve further approximation in the energy equations (for example, in assigning static partial charges to the molecules).^13, 20^ This lack of accuracy is balanced by its much higher speed. Even so, it is unclear that it is suited to the scale of the new ultra-large docking libraries.^7–9^ Recently, a third approach leverages databases of small molecule crystal structures to predict torsion strain using populations of observed angles.^21, 22^ In an influential study, Rarey and colleagues systematically inferred torsion-based ligand strain from the Cambridge Structure Database (CSD)^23^ and also from the Protein Data Bank (PDB).^24^ They compiled histograms of observed dihedral angles for each torsion pattern (the sequence of four atoms defining the dihedral angle, encoded in the SMARTS format).^25^ The torsion patterns were organized hierarchically so that a user can match each torsion pattern in a molecule to the patterns compiled with increasing specificity. They also developed TorsionAnalyzer, which is an interactive graphical tool for strain energy analysis.^26^

Here, we used this statistical approach to address ligand strain energy in docking, focusing exclusively on terms derived from the CSD, which are more accurate. Based on histograms of each torsion pattern, we calculated the torsion energy using a canonical ensemble approach. We converted counts observed in the data sets for different angle measurements for each torsion pattern into Torsion Energy Units (TEU). Since we had a filtering strategy in mind, where torsional strain would either be acceptable or unacceptable but would not be added to a docking score per se, it was unimportant how these TEUs related to docking energies; the two would not be merged. With all counts in the same energy scale, we can compare and add conformational strain energy across different torsion patterns for the entire molecule. We then use the total torsional energy of the molecule, and the maximum individual torsional energy within that molecule, to evaluate the strain of a molecule’s conformation. To determine a threshold for applications, we studied two targets with extensive experimental measurements of docking-predicted ligands, and also 40 systems from the DUD-E benchmark. We find that at certain thresholds, when used as a filter to remove strained molecules, this method can improve docking hit rates, at least retrospectively. The software is relatively fast, capable of calculating strain for half a million molecules on a small cluster in less than 10 minutes, and is mechanically reliable (few molecules fail to return an energy), both of which make it suitable for large library applications. It is openly available to the community at http://tldr.docking.org

## METHODS

### Torsion Library Generation

We represent every sequence of four atoms defining a dihedral angle by a torsion pattern, using the SMARTS line notation.^25^ We adapted the torsion library of Rarey et al.^21, 22^ to build our own library, which has 514 torsion patterns with the same hierarchical organization as the original. Each torsion pattern has a histogram of the observed counts for each possible dihedral angle measurement in the CSD and PDB. For each torsion pattern’s histogram, if the total count is less than 100, we use an approximate approach. The original torsion library allows “tolerances” about each peak in the histogram, where the observed frequency drops below a certain value. Our approximate approach flags any degree difference between a conformation’s dihedral angle and the histogram’s peak that is larger than the maximum tolerance in the original library. If the total count is larger than 100, we convert the histogram frequencies into Torsion Energy Units (TEU) by applying the Boltzmann equation (**SI Figure S1**), which we term the “exact approach”. To avoid infinite energies from zero counts in the histograms, we add the minimum positive count from each histogram to any zero counts. We assume the original measurements from the CSD and PDB exist in a canonical ensemble at 298K, meaning the temperature of the hypothetical ensemble is a tunable parameter in creating the strain energy library. We therefore use TEU instead of kcal/mol to reflect the fact that our energy scale is artificial and based on the databases’ sampling of the hypothetical ensemble. We modeled this unit on Rosetta’s use of Rosetta Energy Units in its scoring function for protein structure conformational energy.^27, 28^

### Workflow of the Software

The current version can handle two types of input files: mol2 and db2 (a format for DOCK 3.7).^29^ With the energy profile for each torsion pattern pre-calculated in the library, we only need to look up the energy estimates for the torsion patterns present in the molecule, not calculate them from scratch. Using the Chem sub-module in the Python module RDKit,^30^ we find all of the torsion patterns present in the molecule based on their SMARTS patterns, calculate their dihedral angles, and then extract the relevant energy estimates from the torsion library. We keep the information matching the most specific (least general) torsion pattern in the library’s hierarchy. Then we can sum the energy estimates for all of the torsion patterns to get the estimate for the molecule’s conformation, or simply find the torsion patterns with energy estimates above a desired cutoff (**Figure 1**).

**Figure 1.**
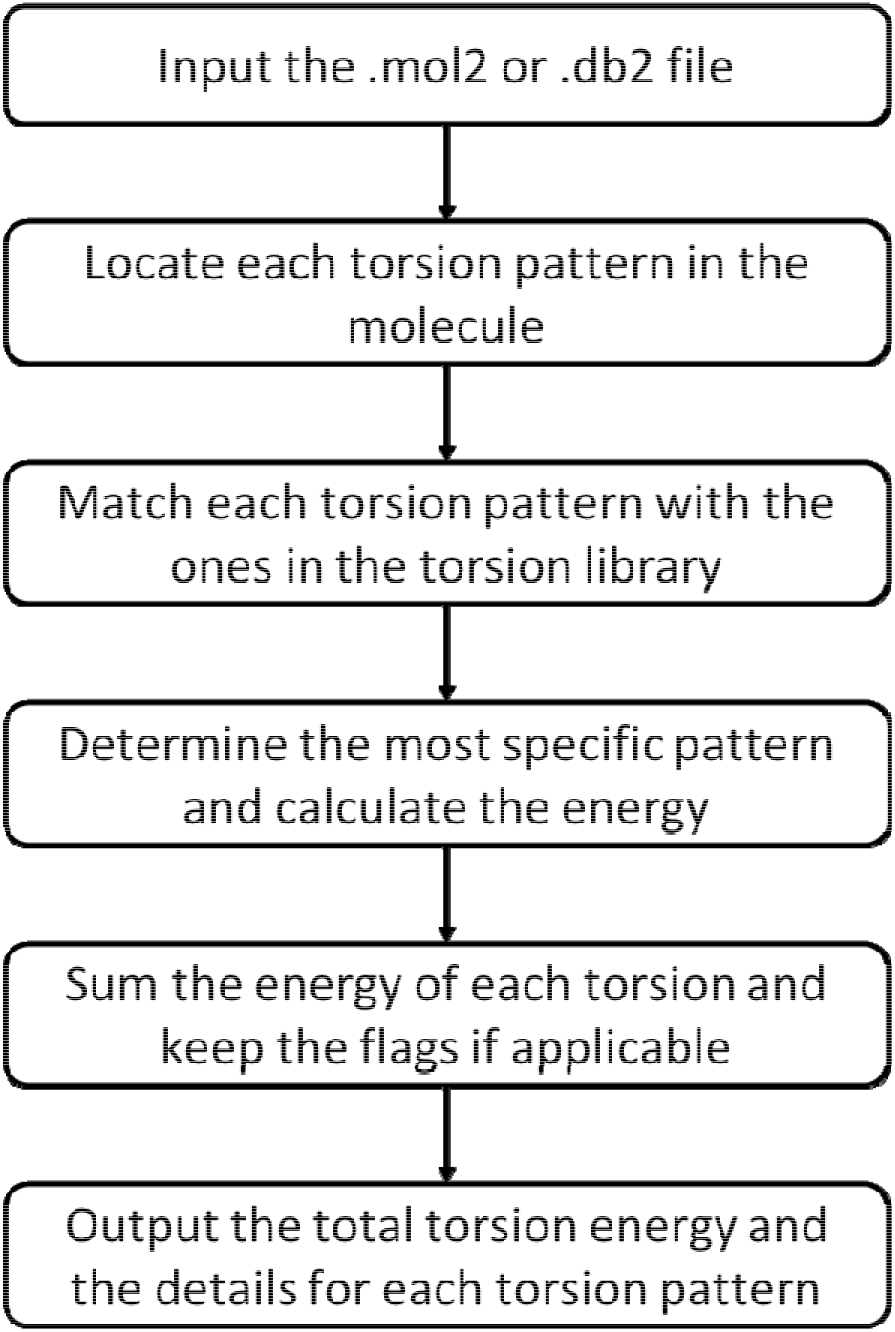
Flowchart for strain energy filtering. The program first locates each torsion pattern in the molecule and calculates its dihedral angle. It then matches each torsion pattern in the molecule with the patterns in the torsion library. There will be multiple such matches, since the torsion library contains a hierarchy of patterns. For each match, the program calculates the energy for the observed dihedral angle and determines any flags. For each torsion pattern in the molecule, it keeps only the information from the most specific torsion pattern rule from the library. Ultimately, the program reports the estimated energy for each torsion pattern, the sum for all the patterns in the molecule, and any flagged patterns.

### DUD-E Benchmark Docking

The full list of DUD-E systems and the corresponding PDBIDs used are listed in **Table S3**. For docking, the crystallographic ligands were used to generate matching spheres for each target. To prepare the protein in docking, we pre-calculated energy grids for an AMBER-based van der Waals potential,^31^ a Poisson–Boltzmann electrostatic potential, using QNIFFT,^32, 33^ and for ligand desolvation using an adapted Generalized-Born approach.^34^ We used DOCK3.7^29^ to dock all the hits and decoys for each DUD-E target.

## RESULTS

### D4 Dopamine Receptor Case Study

We began by investigating the effects of ligand strain on the experimental results of large library docking campaign against the D4 dopamine receptor.^7^ Here, 549 docking-ranked compounds, across a wide range of ranks, were tested in the same assay in the same lab, providing an unusually large set of comparable experiments for a diverse set of compounds. We collected 256 of these molecules that had high DOCK3.7 ranks, with scores less (better) than −60 kcal/mol.^7^ Of these, 62 were found experimentally to bind, with EC_50_ values ranging from 6 μM to 180 pM, while the other 194 were docking decoys (i.e., high-ranking molecules that did not bind on experimental testing). In the calculation that led to these molecules, no strain energy filter was applied. We can calculate the torsion-based ligand strain for each of these experimental binders and non-binders, comparing the docked conformation of the molecules to that of their ground state (by torsion strain) (**Table S1**). If it is true that the decoys tend to be more strained than the binders, then applying a strain filter should remove more of the experimental non-binders than the true binders, and correspondingly the hit rates— the number-active/number-tested—should rise. These values can be calculated at different strain thresholds (**Figure 2**). Here, “Total” refers to the total strain energy of a compound, adding up all torsions, while “Single” means the maximum strain energy of any individual torsion within the molecule. We deem a molecule’s docked pose “strained” if its total or maximum individual torsion strain is greater than the threshold.

**Figure 2.**
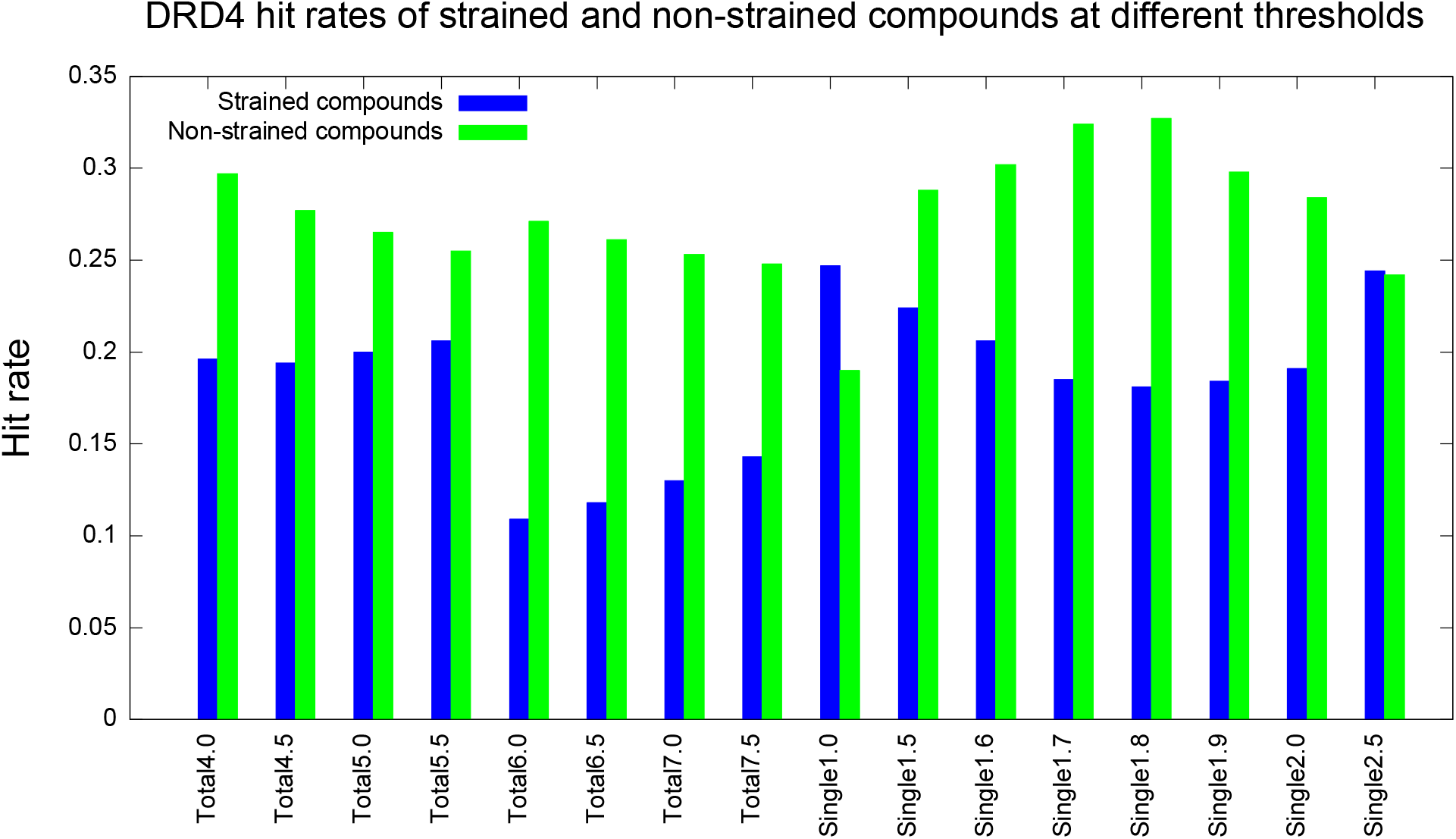
D4 receptor hit rates of strained (blue) and non-strained (green) compounds at different thresholds, measured in TEU. For the total energy category, 6.0 TEU appears to be a good threshold. For the maximum single energy category, 1.8 TEU appears to be a good choice.

A satisfactory strain energy correction should maximize the hit rate of non-strained compounds while minimizing that of the strained ones, thus maximizing the difference between them. The hit rate of non-strained is the number of true positive (true hits and correctly labeled as non-strained) divided by the number of non-strained compounds, while the hit rate of strained is the number of false negative (true hits but wrongly labeled as strained) divided by the number of strained compounds. In the case study of the D4 receptor, we maximize the difference in hit rates using a total strain energy threshold of 6.0 TEU and a maximum single strain energy threshold of 1.8 TEU.

Inspection of the compounds filtered out or retained affords insight into this process (**Figure 3)**. High-ranking decoys like ZINC000067715288 and ZINC000666614106 break planarity about an amide bond, with an angle of about 30 degrees or more, imparting a single energy strain of 4.0 and 5.6 TEUs, respectively, and a Total strain of 10.4 and 5.8 TEUs, respectively. Molecules like ZINC000377646411 and ZINC000248536951 have strained torsions that break conjugation of an exocyclic group with an aromatic ring. These strained torsions allow these molecules to make favorable interactions with the receptor that they otherwise could not (**Figure 3**). Conversely, for high-ranking binders like ZINC000480496068 and ZINC000155719879 their torsional violations never exceed 1.2 TEUs for any single angle, nor do they add up to more than 4.2 in Total.

**Figure 3.**
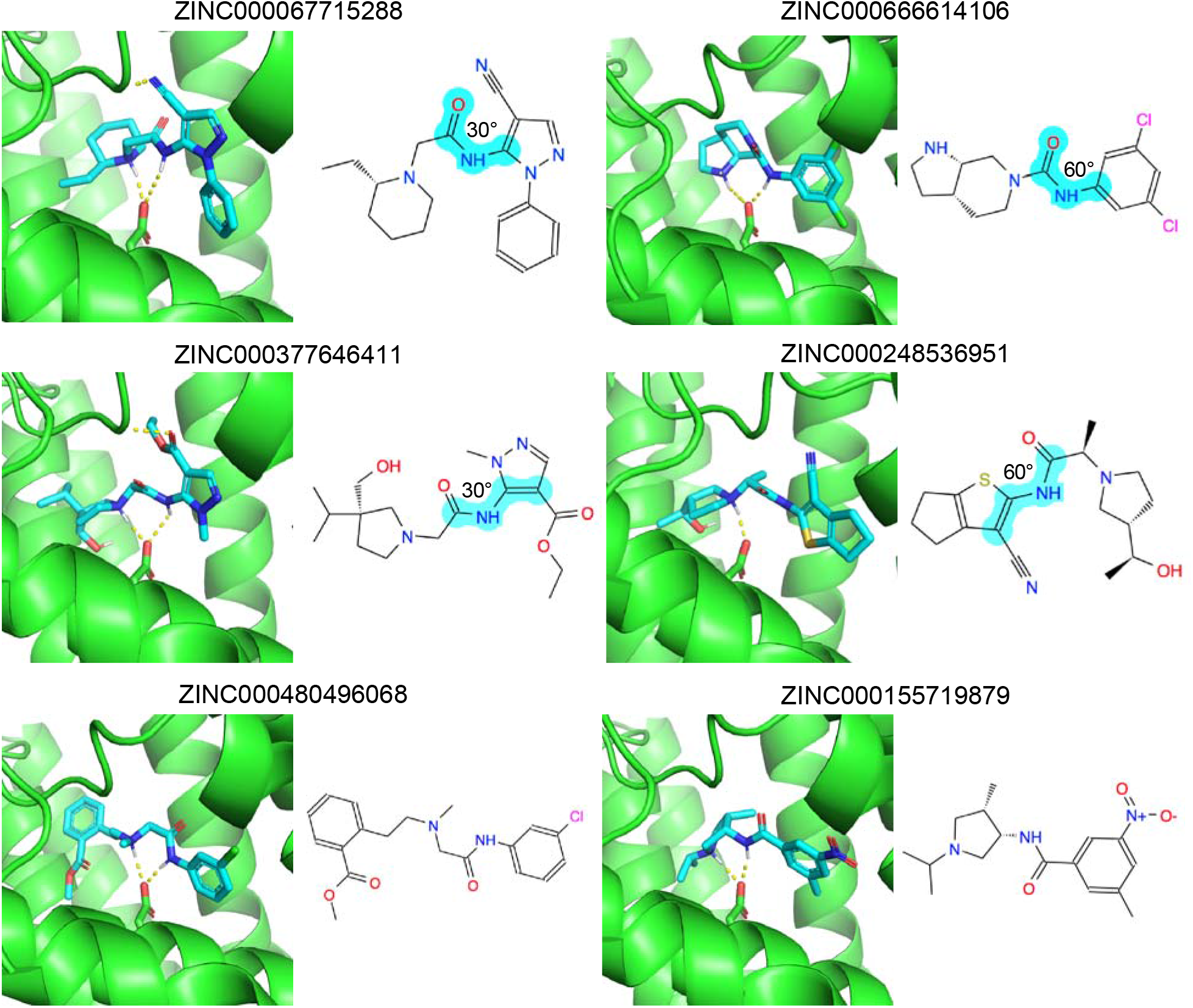
Docking poses of strained decoys (top four) and unstrained binders (bottom two) in DRD4. 2D images of the molecules are also shown. The strained torsions are indicated in cyan and labeled with its degrees out of optimum.

We looked at the effects of strain energy filtering on the docking prioritized D4 receptor compounds, and on the true ligands that emerged from them, using a total strain of 6.0 TEU and maximum single strain of 1.8 TEU (**Table 1**). Using the total strain filter alone (columns 3 and 4), the hit rates of strained and non-strained compounds—i.e., above and below our threshold— are 0.109 and 0.271, respectively. A two-sample proportion Z-test indicates that this difference is significant (p-value 0.010). Using the maximum single strain filter (columns 5 and 6), the hit rates of strained and non-strained compounds are 0.181 and 0.327, respectively, which is also a significant difference (p-value 0.004). The p-value for the hit rate comparison with and without the maximum single strain filter (columns 6 and 2) is 0.048, which is significant. On the other hand, the p-value for the hit rate comparison with and without the total strain filter (columns 4 and 2) is 0.236. While this p-value is not significant, an even more stringent criteria where both thresholds are used can be imagined.

**Table 1.** DRD4 hit rates before and after strain filtering at two chosen thresholds.

| | All (no strain filters) | Strained after applying strain filter (total strain ≥ 6 TEU) | Non-strained (total strain < 6 TEU) | Strained after applying strain filter (max single strain ≥ 1.8 TEU) | Non-strained (max single strain < 1.8 TEU) |
| --- | --- | --- | --- | --- | --- |
| Number of hits | 62 | 5 | 57 | 27 | 35 |
| Sample size | 256 | 46 | 210 | 149 | 107 |
| Hit rate | 0.242 | 0.109 | 0.271 | 0.181 | 0.327 |

Although each threshold is the best choice in its category (with this level of granularity in testing thresholds), the outcomes are different. At total strain 6.0 TEU, only 46 compounds are filtered out vs 149 at maximum single strain 1.8 TEU; the percentage of remaining compounds are 82% versus 42%, respectively. This difference suggests that if the investigator prefers to keep as many hits as possible (conservative in deeming a molecule “strained”), they should use the total strain threshold. To maximize stringency, ensuring that most compounds will be “unstrained”, they should use the maximum single strain threshold.

### Case Study of AmpC β-Lactamase

We applied the same calculations and analyses on the AmpC β-lactamase dataset, another target for which we have a substantial number of true actives and experimentally measured decoys from a previous large-scale docking campaign. Of 44 experimentally-tested molecules with DOCK score less than −60,^7^ 5 were true inhibitors (39 were high-ranking decoys) (**Table S2, Figure 4**).

**Figure 4.**
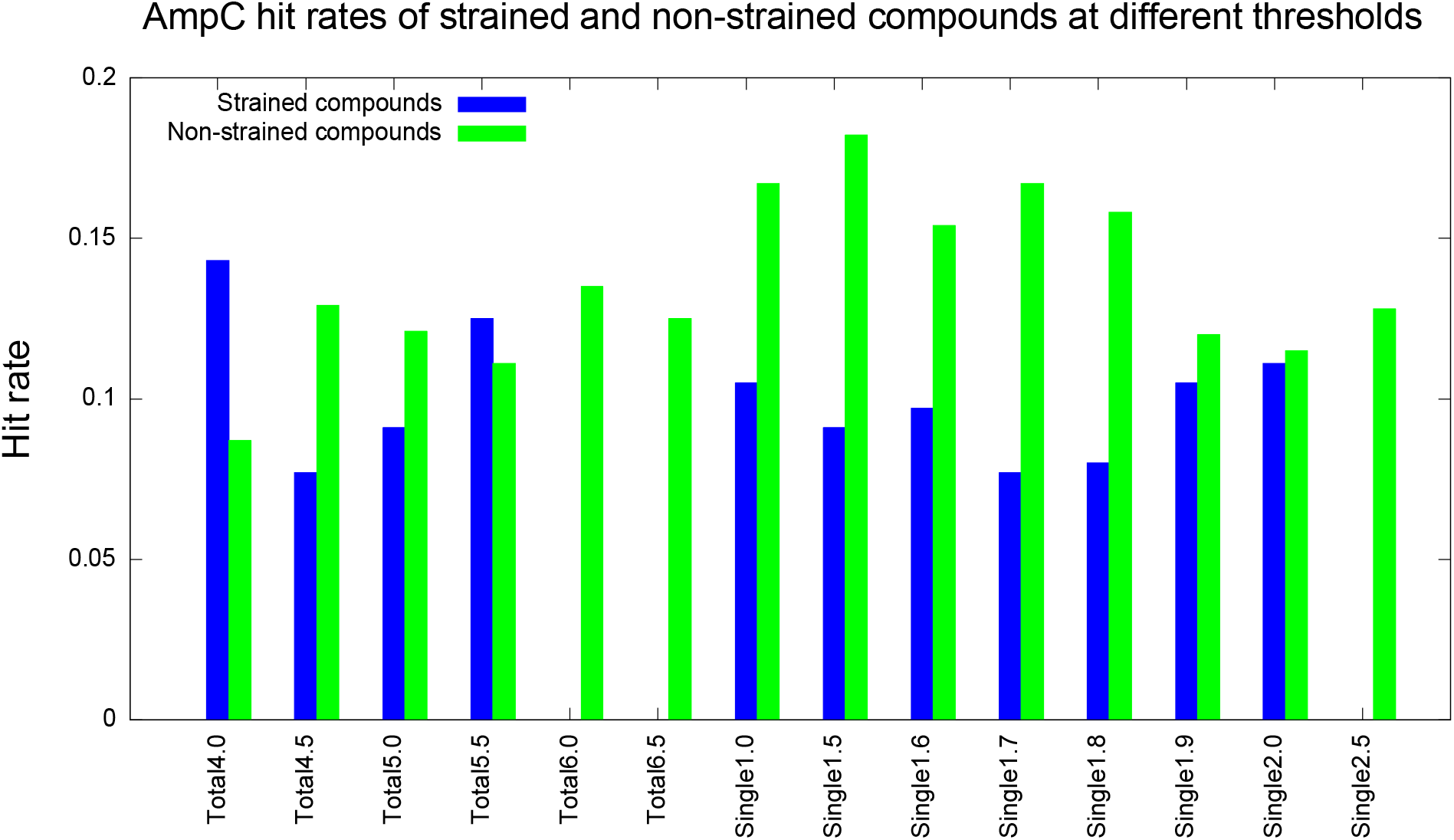
AmpC hit rates of strained (blue) and non-strained (green) compounds at different thresholds, measured in TEU. For the total energy category, 6.0 TEU is a good choice of threshold. For the maximum single energy category, 1.7 TEU is a good choice.

As with the D4 case, we maximize the difference in hit rates between strained and non-strained molecules for the AmpC molecules. Here, the maximum difference between the two comes with a total strain energy threshold of 6.0 TEU and a maximum single strain energy threshold of 1.7 TEU (**Table 2**). Using the total strain energy filter (columns 3 and 4), the hit rates of strained and non-strained compounds are 0.000 and 0.135, respectively (i.e., none of the true ligands are found to be strained). Using the maximum single strain energy filter (columns 5 and 6), the hit rates of strained and non-strained compounds are 0.077 and 0.167, respectively. The two-sample proportion Z-test failed to show statistical significance for these differences, as the total sample size of 44 was too small. Still, overall, we observed similar results for AmpC as we did for D4. At total strain 6.0 TEU, only 7 compounds are filtered out compared to 26 at single maximum strain 1.7 TEU. The percentage of remaining compounds are 84% versus 41%. As we will see, this trend continues when we turn to the DUD-E benchmarks.

**Table 2.** AmpC hit rates before and after strain filtering at two chosen thresholds.

| | All (no strain filters) | Strained after applying strain filter (total strain $\geq 6$ TEU) | Non-strained (total strain $< 6$ TEU) | Strained after applying strain filter (max single strain $\geq 1.7$ TEU) | Non-strained (max single strain $< 1.7$ TEU) |
| --- | --- | --- | --- | --- | --- |
| Number of hits | 5 | 0 | 5 | 2 | 3 |
| Sample size | 44 | 7 | 37 | 26 | 18 |
| Hit rate | 0.114 | 0.000 | 0.135 | 0.077 | 0.167 |

Here too, we can inspect the conformations adopted by the compounds filtered out by strain (**Figure 5)**. High-ranking decoys like ZINC000188308020 and ZINC000261754704 break planarity about an amide bond, with an angle of about 30 degrees, imparting a single energy strain of 2.1 and 2.1 TEUs, respectively, and a Total strain of 6.8 and 6.6 TEUs, respectively. Molecules like ZINC000317738578 and ZINC000274934818 have strained sulfonamide with nitrogen in a ring. These strained torsions allow these molecules to make favorable interactions with the enzyme that they otherwise could not (**Figure 5**). Conversely, for high-ranking binders like ZINC000184991516 and ZINC000547933290, their torsional violations never exceed 1.7 TEUs for any single angle, nor do they add up to more than 4.5 in Total; nevertheless, they make favorable fits with β-lactamase, comfortably placing a phenolate in the oxyanion hole, and hydrogen bonding with the key recognition Asn152 of the enzyme.

**Figure 5.**
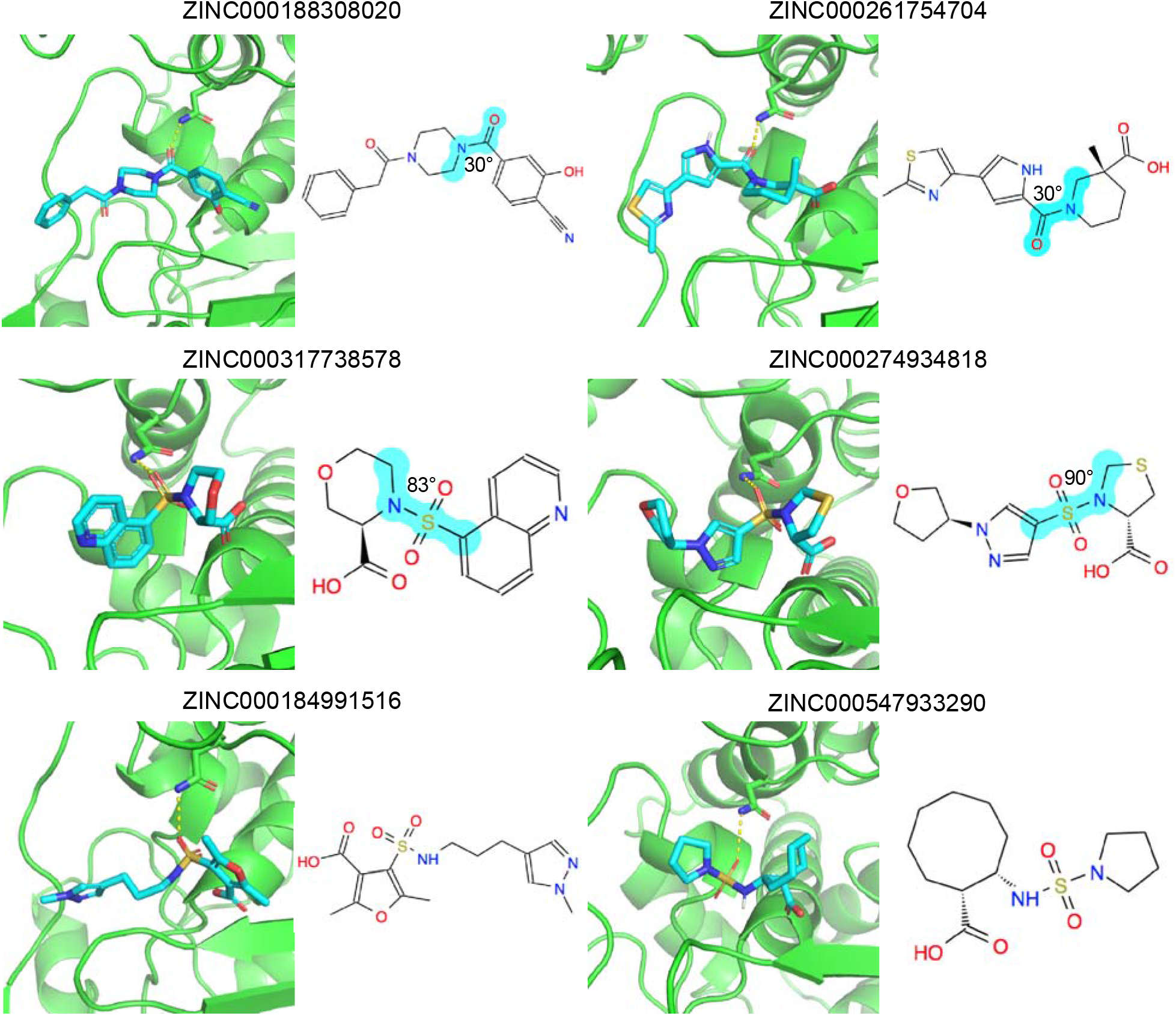
Docking poses of strained decoys (top four) and unstrained binders (bottom two) in AmpC. 2D images of the molecules are also shown. The strained torsions are indicated in cyan and labeled with its degrees out of optimum.

### DUD-E Benchmark Tests

After the case studies of the D4 receptor and AmpC, we investigated performance against targets from the DUD-E benchmark.^35^ Here, the decoys are not, as in D4 and AmpC, experimentally measured non-binders, but are based on topological differences from the known ligands. The DUD-E database is a widely-used benchmark to test docking.^35, 36^ It includes 102 targets with an average of 224 ligands each, and 50 property-matched decoys for each ligand. Compared to AmpC and the D4 receptor, DUD-E targets have the disadvantage of depending on presumed non-binders vs the experimentally determined non-binders afforded to us by AmpC and D4. This is balanced by the many DUD-E targets, spanning a wide range of chemotypes, and their wide use in the field.

Accordingly, we docked the ligands and decoys for 40 DUD-E systems against their targets, and calculated the adjusted LogAUC^37^ for integrated ligand enrichment before and after strain filtering (LogAUC measures the area under the enrichment curve over the database docked, weighting different order-of-magnitude regions equally, with the range 0.1 to 1% of molecules docked weighted the same as 1 to 10% and 10 to 100%, thereby up-weighting the high-ranking region; the method is adjusted by subtracting the LogAUC expected at random). We used the change in adjusted LogAUC (ΔLogAUC), the LogAUC after strain filtering minus the LogAUC before strain filtering, to measure method performance: a positive ΔLogAUC indicates improvement by filtering for molecular strain.

We measured the effect of different strain energy filter values on integrated enrichment on the DUD-E benchmark (**Figure 6**), with the left Y-axis the ΔLogAUC at different thresholds (blue bars). As we decrease the value of the strain energy below which molecules are filtered out (stringency increases), ΔLogAUC increases. Said another way, as one filters out compounds with greater-and-greater stringency (lower and lower strain energy allowed), enrichment improves. For instance, filtering out any compound with total strain greater than 4 TEU improves ΔLogAUC more than only filtering out compounds with total strain greater than 8 TEU. When we filter on the single largest torsion energy, the effect is less monotonic, with 1.5 TEU returning the largest ΔLogAUC, followed by 1.0 and 1.8 TEU. These effects must be balanced against the number of compounds remaining after strain filtering. At thresholds of 1.0 TEU and 1.5 TEU for the single torsional strain, the percentage of total remaining compounds fall to only 9.6% and 22.1%, respectively (**SI Figure S2)**. Accordingly, we preferred a maximum single strain threshold of 1.8 TEU over 1.0 or 1.5 TEU.

**Figure 6.**
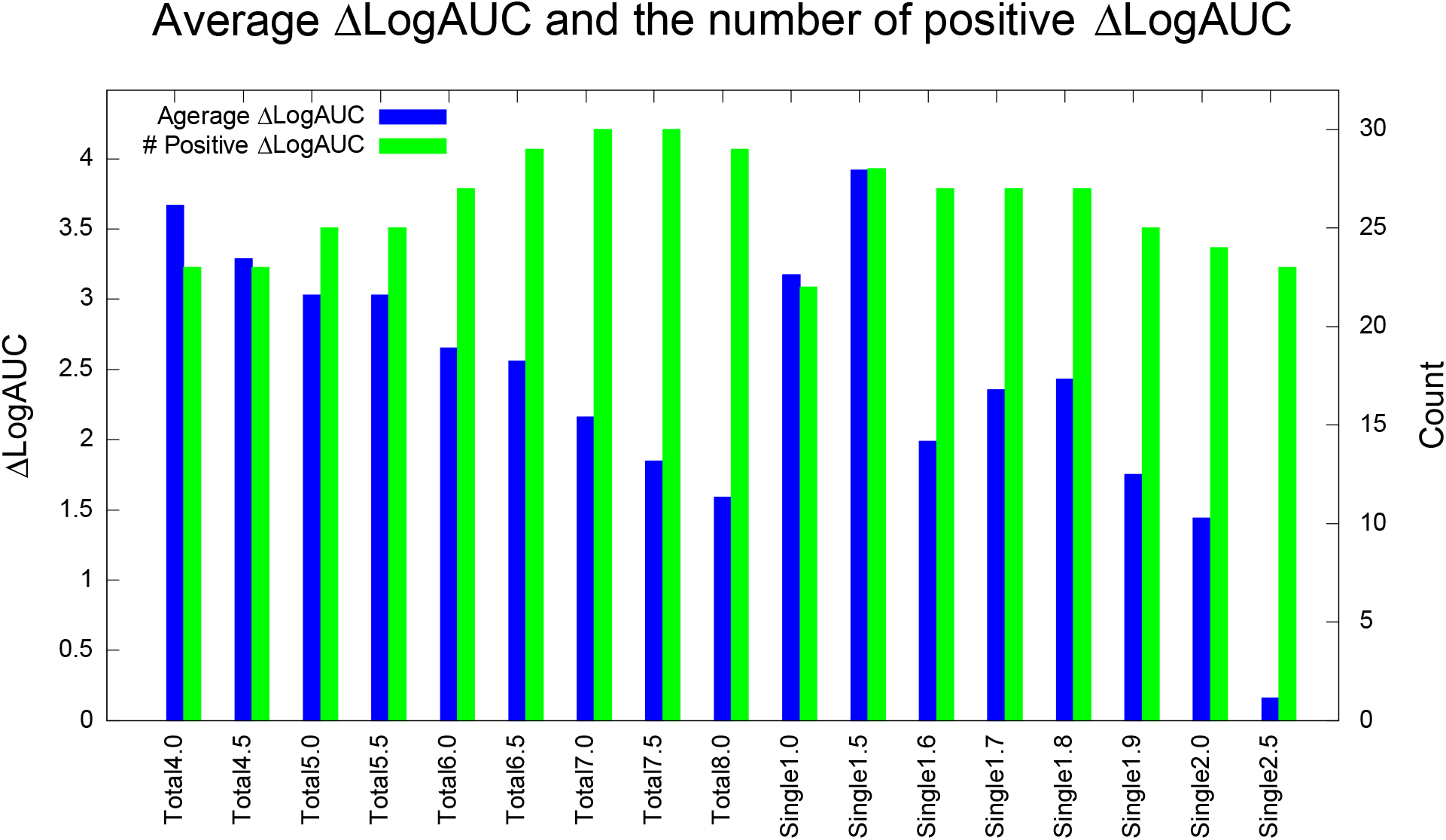
DUD-E benchmark tests at different thresholds (in TEU). Blue is the average ΔLogAUC and green is the number of positive ΔLogAUC among 40 systems.

The effect of the strain energy filters may be also evaluated by the number of systems where integrated enrichment improved (green bars in **Figure 6**, right Y-axis, over 40 systems). Here the trend was almost opposite that for the average ΔLogAUC—as stringency diminished, the total number of systems that improved increased (even though the overall average dropped). Filtering by total strain energy at 7.0 and 7.5 TEU saw the highest number of total systems showing improvement, with 30 targets overall—75% of the systems evaluated—showing improved enrichment. Filtering by the single larger torsional energy in the molecules, 28 systems improved at 1.5 TEU, while filtering at 1.6, 1.7, and 1.8 TEU all led to improvements in 27 systems. If we take total 7.0 TEU and maximum single strain 1.8 TEU as filters, an average of 71.4% and 37.1% compounds survive these filters, respectively. This is consistent with what we observed with the D4 receptor and with AmpC β-lactamase: filtering by total strain energy retains more compounds, while filtering by single maximum torsional strain is more stringent. How best to use and combine these filters may depend on context. In large-scale docking, for instance, one is often confronted with an embarrassment of riches, with too many high-scoring molecules to choose; here, the more stringent maximum single torsional filter may be the better choice. Strategies that combine the two criteria may also be imagined.

The results may be broken down by the individual protein in the 40-target benchmark, for simplicity just using a total strain of 7.0 and a maximum single torsional strain of 1.8 TEU (**Figure 7**). Using the maximum strain filter, 30 targets had improved LogAUC, while at the maximum single torsional strain 27 targets did. For some targets like Renin (RENI) and Tryptase beta-1 (TRYB1), strain filtering greatly boosted in the LogAUC. The ligands of these targets are usually large with multiple rotatable bonds. In **SI Figure S3**, we plot ΔLogAUC vs average number of ligand dihedral angles for all the targets, observing a positive correlation between the total strain and the number of ligand and decoy rotatable bonds.

**Figure 7.**
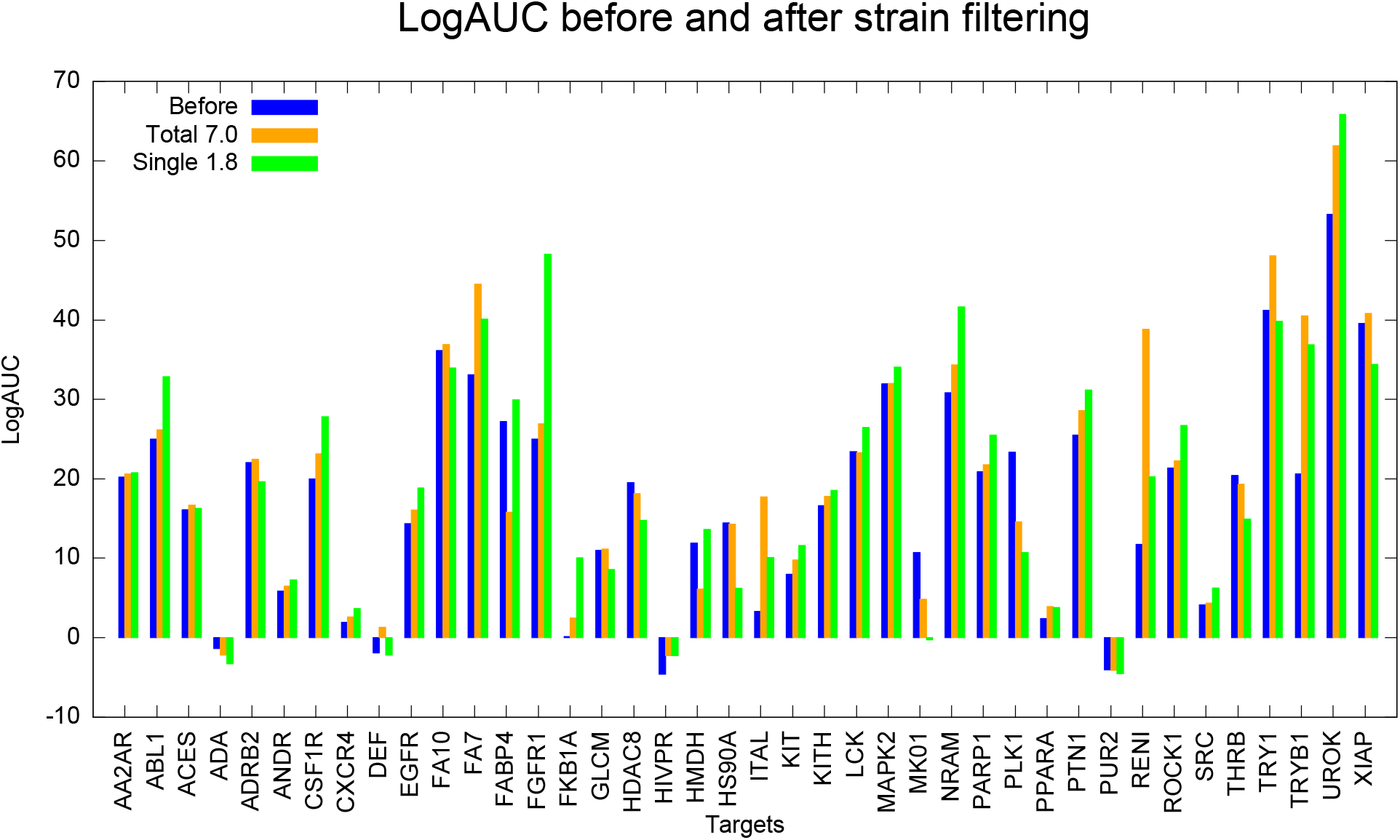
LogAUC before and after strain filtering of 40 systems. Blue is the data before strain filtering. 30 targets improved at the total strain threshold of 7.0 TEU (orange), while 27 improved at the maximum single strain threshold of 1.8 TEU (green).

As with the D4 and AmpC case studies, we can inspect the conformations adopted by the compounds filtered out by strain for the DUD-E targets (**Figure 8)**. High-ranking decoys like C26299839 and C40966046 break planarity about an amide bond, with an angle of about 30 degrees or more, imparting a single energy strain of 2.1 and 3.4 TEUs, respectively, and a Total strain of 8.4 and 9.5 TEUs, respectively. Molecules like C38700416 and C09344195 have strained torsions that disrupt conjugation of an extra-cyclic group with an aromatic ring. These strained torsions allow these molecules to make favorable interactions with the targets that they otherwise could not (**Figure 8**).

**Figure 8.**
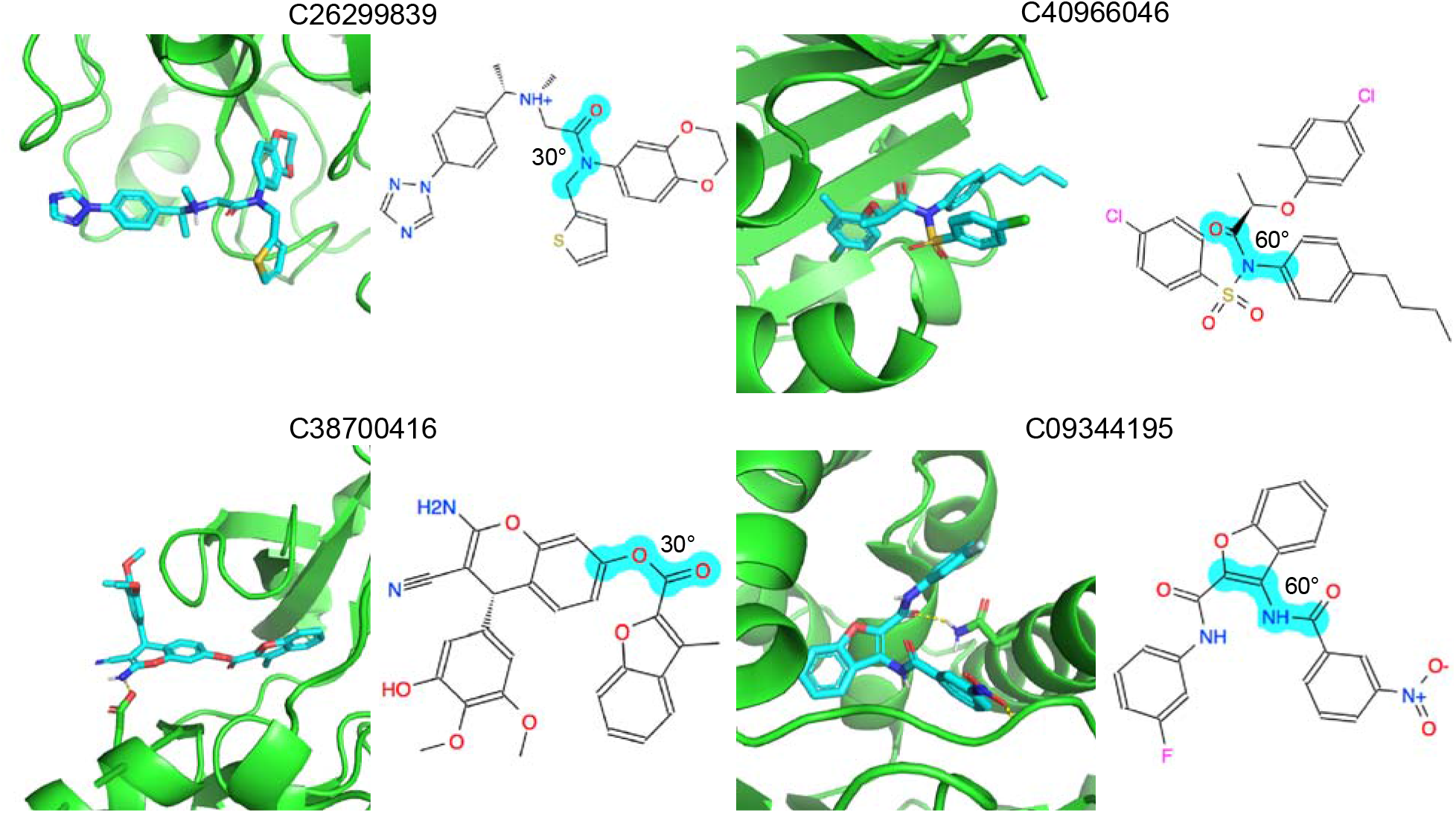
Docking poses of four strained decoys: C26299839 in FA10, C40966046 in ITAL, C38700416 in ABL1, and C09344195 in HS90A. 2D images of the molecules are also shown. The strained torsions are indicated in cyan and labeled with its degrees out of optimum.

### Speed and Robustness

To investigate speed and robustness, we calculated strain energies on three sets of 500,000 compounds each. On a desktop of i5-8400 CPU @ 2.80GHz, 16 GB RAM and CentOS Linux 7.6.1810, it took an average of 319 minutes to calculate the strain energy for the half million compounds, or less than 0.04 second for each compound. This is fast enough to be used as a post-docking filter. We further did a parallel test on our cluster, splitting the 500,000 compounds into 100 jobs, each with 5,000 compounds. On Intel Xeon Silver 4210 CPU @ 2.20GHz, the running time of each job ranged from 301 to 463 seconds, with an average of 390. With even a small cluster of 100 cores, this strain energy calculation can be conducted for half million molecules in eight minutes. While this project has focused on using strain as a filter for docking results at the end of a large library campaign, the speed of this calculation suggests it may be possible to use for library generation, even for the >1 billion molecule ultra-large libraries. Speaking to robustness, in the three tests the failure rate was <5 per million compounds; almost all the provided compounds could be properly processed. The ones that failed are usually compounds without any dihedrals, like benzopyrene.

## DISCUSSION

Applying a strain energy filter to docked compounds improves docking hit rates and integrated enrichment for most targets, at least retrospectively. This is true in cases where both ligands and “decoys” derive from molecules from prospective docking screens that have been shown to bind or not bind by experiment, respectively (as with AmpC and the D4 dopamine receptor), and for cases where the putative non-binders are property-matched decoys, of the sort found in the DUD-E benchmarks. The reason that this strain filter improves hit-rates and enrichments is because the decoy molecules, both experimental and calculated, tend to find their best fitting conformations from among those that are relatively strained, compared to the known ligands that can more often adopt high-complementary conformations that are relatively unstrained.

In their seminal paper on conformational strain in docking, Tirado-Rives and Jorgensen argued that the errors in both the strain calculations and in overall docking scores were so high—over the 5 kcal/mol or so that defined the range of docking binding energies—as to make even rank-ordering a dubious proposition.^1^ While this likely remains true, our results may be reconciled with theirs by distinguishing between rank ordering, the focus of their study, and enrichment, the goal of this study. To improve docking enrichment, a strain energy does not need to be even included in the overall docking score, it can be used to filter out molecules that only fit well because they adopt strained conformations. Because these fall more often among decoys than the true ligands this filter may typically improve docking results.

Several caveats merit airing. Most importantly, this study is retrospective, and the true value of a new docking tool will often only emerge in prospective studies. Mechanically, we calculated LogAUC using the compounds that remained after strain filtering, while those filtered-out were not included. Another way of calculating LogAUC would be to include all the compounds—both ligands and decoys—whether filtered or not. We chose the former to be consistent with the real practice of docking, where one only tests compounds that survive various filters and are prioritized by high ranks. From the standpoint of whether we should be using filters at all, admittedly a strain filter is only needed because library molecules have been calculated in strained conformations. If these were recognized before docking, the need for this filter would disappear. Indeed, Omega (OpenEye Software, Santa Fe), which we use to pre-calculate the flexibase for docking,^38^ and which is among the premier methods in the field, offers the opportunity to do just that, using essentially the same statistical potential on which we draw here. In our hands, implementing this at the time of conformation generation under-sampled ligand-competent conformations, reducing retrospective hit rates. It may well be possible to overcome this in subsequent implementations. One could further imagine integrating strain energies directly into docking scores. This would have the advantage of being truer to the biophysics, but, as pointed out by Tirado-Rives & Jorgensen,^1^ would struggle to balance the other docking confounds. Our own view is that a strain energy filter, outside of conformer generation and sampling, may have an advantage in docking, as it is unhobbled by outside dependencies. Finally, one could easily imagine implementing ligand conformational strain directly into the docking library, where it could be applied at the time of docking, and not used as a post-docking filter. This would bring considerable advantages, and could conceivably be done with methods similar to that used here. Whether this will be pragmatic for ultra-large libraries demands further exploration, outside of the scope of this study.

These caveats should not obscure the principle observations of this study: despite ongoing difficulties with rank-ordering in docking,^1^ a strain energy filter can improve docking enrichment—the first and principal goal of library screening. Especially in the era of ultra-large libraries, where one suffers from problems of abundance, stringency filters like strain energy can be a great advantage by removing lower likelihood candidates—other such filters can also be imagined. Drawing on a database torsion angle approach,^21, 22^ the filter we describe is fast and mechanically reliable, able to treat millions of molecules in minutes; even from a highly diverse library, only 1 in 200,000 molecules fails to have its torsions matched and an energy calculated. Applying such a filter to docked conformations may increase prospective hit rates by eliminating non-binders that rank well only by adoption of high-energy conformations. The software is openly available to the community (http://tldr.docking.org).

## ASSOCIATED CONTENT

### Supporting Information

An example of torsion energy profile of a SMARTS pattern (Figure S1); average percentage of remained compounds after strain filtering at different conditions (Figure S2); correlation between ΔLogAUC and the ligands’ average number of dihedrals at the thresholds total strain 7.0 TEU (Figure S3); ligands docking score and strain energy in D4 receptor (Table S1); ligands docking score and strain energy in AmpC (Table S2); DUD-E LogAUC results before and after strain filtering at different thresholds (Table S3)

## AUTHOR INFORMATION

### Notes

The authors declare no competing financial interest.

## ACKNOWLEDGEMENTS

This study was supported by R35GM122481 (to BKS). We thank the contributors for RDKit, on which we drew heavily, and OpenEye scientific for Omega. We thank members of the Shoichet Lab for testing the software and for helpful discussion, and Stefan Gahbauer and Brian Bender for reading this manuscript.

## ABBREVIATIONS

AA2AR: Adenosine A2a receptor
ABL1: Tyrosine-protein kinase ABL
ACES: Acetylcholinesterase
ADA: Adenosine deaminase
ADRB2: Beta-2 adrenergic receptor
ANDR: Androgen Receptor
CSF1R: Macrophage colony stimulating factor receptor
CXCR4: C-X-C chemokine receptor type 4
DEF: Peptide deformylase
EGFR: Epidermal growth factor receptor erbB1
FA10: Coagulation factor X
FA7: Coagulation factor VII
FABP4: Fatty acid binding protein adipocyte
FGFR1: Fibroblast growth factor receptor 1
FKB1A: FK506-binding protein 1A
GLCM: Beta-glucocerebrosidase
HDAC8: Histone deacetylase 8
HIVPR: Human immunodeficiency virus type 1 protease
HMDH: HMG-CoA reductase
HS90A: Heat shock protein HSP 90-alpha
ITAL: Leukocyte adhesion glycoprotein LFA-1 alpha
KIT: Stem cell growth factor receptor
KITH: Thymidine kinase
LCK: Tyrosine-protein kinase LCK
MAPK2: MAP kinase-activated protein kinase 2
MK01: MAP kinase ERK2
NRAM: Neuraminidase
PARP1: Poly [ADP-ribose] polymerase-1
PLK1: Serine/threonine-protein kinase PLK1
PPARA: Peroxisome proliferator-activated receptor alpha
PTN1: Protein-tyrosine phosphatase 1B
PUR2: GAR transformylase
RENI: Renin
ROCK1: Rho-associated protein kinase 1
SRC: Tyrosine-protein kinase SRC
THRB: Thrombin
TRY1: Trypsin I
TRYB1: Tryptase beta-1
UROK: Urokinase-type plasminogen activator
XIAP: Inhibitor of apoptosis protein

## Table of Contents Graphic

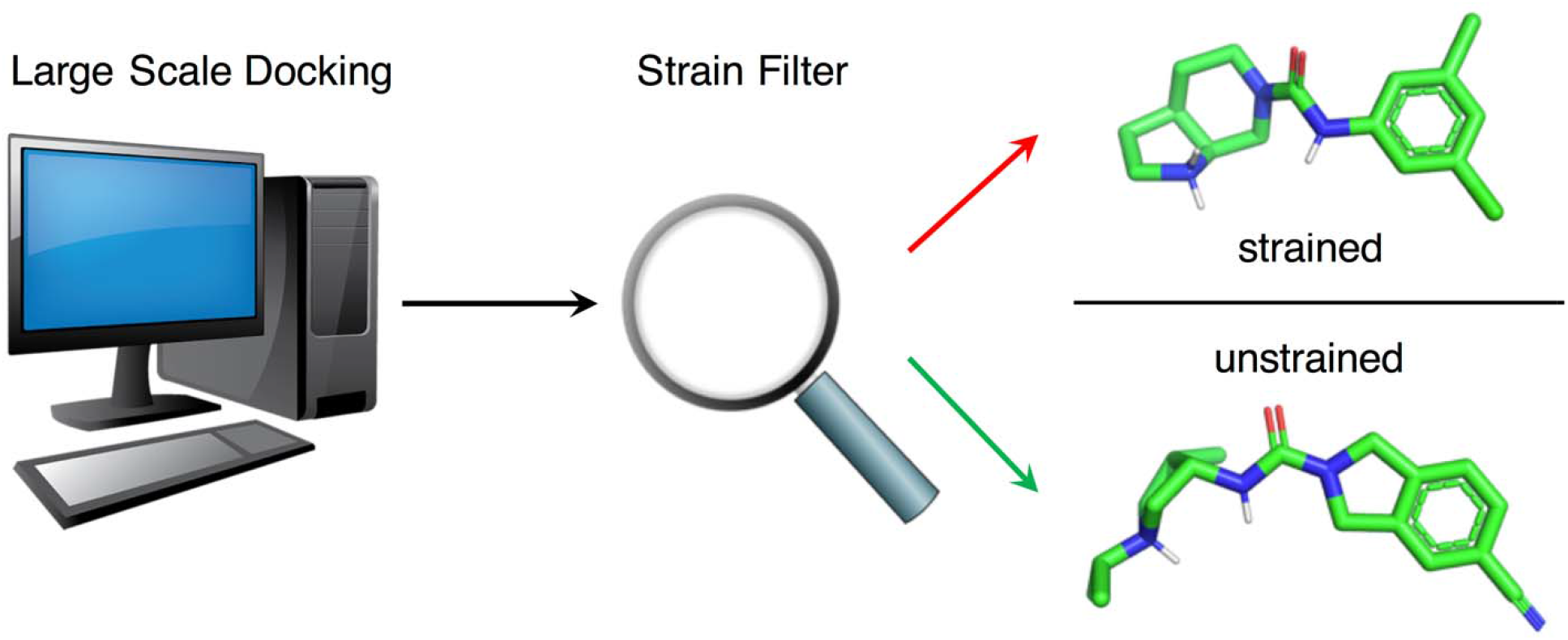

## REFERENCES

1. Tirado-Rives, J.; Jorgensen, W. L. J. J. o. m. c., Contribution of conformer focusing to the uncertainty in predicting free energies for protein− ligand binding. 2006, 49, 5880–5884.

2. Cecchini, M.; Kolb, P.; Majeux, N.; Caflisch, A., Automated docking of highly flexible ligands by genetic algorithms: a critical assessment. J Comput Chem 2004, 25, 412–22.

3. Friesner, R. A.; Banks, J. L.; Murphy, R. B.; Halgren, T. A.; Klicic, J. J.; Mainz, D. T.; Repasky, M. P.; Knoll, E. H.; Shelley, M.; Perry, J. K.; Shaw, D. E.; Francis, P.; Shenkin, P. S., Glide: a new approach for rapid, accurate docking and scoring. 1. Method and assessment of docking accuracy. J Med Chem 2004, 47, 1739–49.

4. Murphy, R. B.; Repasky, M. P.; Greenwood, J. R.; Tubert-Brohman, I.; Jerome, S.; Annabhimoju, R.; Boyles, N. A.; Schmitz, C. D.; Abel, R.; Farid, R.; Friesner, R. A., WScore: A Flexible and Accurate Treatment of Explicit Water Molecules in Ligand-Receptor Docking. J Med Chem 2016, 59, 4364–84.

5. Lam, P. C.; Abagyan, R.; Totrov, M., Ligand-biased ensemble receptor docking (LigBEnD): a hybrid ligand/receptor structure-based approach. J Comput Aided Mol Des 2018, 32, 187–198.

6. Mobley, D. L.; Dill, K. A., Binding of small-molecule ligands to proteins:“what you see” is not always “what you get”. Structure 2009, 17, 489–498.

7. Lyu, J.; Wang, S.; Balius, T. E.; Singh, I.; Levit, A.; Moroz, Y. S.; O’Meara, M. J.; Che, T.; Algaa, E.; Tolmachova, K., Ultra-large library docking for discovering new chemotypes. Nature 2019, 566, 224–229.

8. Stein, R. M.; Kang, H. J.; McCorvy, J. D.; Glatfelter, G. C.; Jones, A. J.; Che, T.; Slocum, S.; Huang, X. P.; Savych, O.; Moroz, Y. S.; Stauch, B.; Johansson, L. C.; Cherezov, V.; Kenakin, T.; Irwin, J. J.; Shoichet, B. K.; Roth, B. L.; Dubocovich, M. L., Virtual discovery of melatonin receptor ligands to modulate circadian rhythms. Nature 2020, 579, 609–614.

9. Gorgulla, C.; Boeszoermenyi, A.; Wang, Z. F.; Fischer, P. D.; Coote, P. W.; Padmanabha Das, K. M.; Malets, Y. S.; Radchenko, D. S.; Moroz, Y. S.; Scott, D. A.; Fackeldey, K.; Hoffmann, M.; Iavniuk, I.; Wagner, G.; Arthanari, H., An open-source drug discovery platform enables ultra-large virtual screens. Nature 2020, 580, 663–668.

10. Allen, F. H.; Harris, S. E.; Taylor, R., Comparison of conformer distributions in the crystalline state with conformational energies calculated by ab initio techniques. Journal of computer-aided molecular design 1996, 10, 247–254.

11. Butler, K. T.; Luque, F. J.; Barril, X., Toward accurate relative energy predictions of the bioactive conformation of drugs. Journal of computational chemistry 2009, 30, 601–610.

12. Wei, W.; Champion, C.; Barigye, S. J.; Liu, Z.; Labute, P.; Moitessier, N., Use of Extended-Hückel Descriptors for Rapid and Accurate Predic ons of Conjugated Torsional Energy Barriers. Journal of Chemical Information Modeling 2020, 60, 3534–3545.

13. Sellers, B. D.; James, N. C.; Gobbi, A., A Comparison of Quantum and Molecular Mechanical Methods to Estimate Strain Energy in Druglike Fragments. J Chem Inf Model 2017, 57, 1265–1275.

14. Rai, B. K.; Sresht, V.; Yang, Q.; Unwalla, R.; Tu, M.; Mathiowetz, A. M.; Bakken, G. A., Comprehensive Assessment of Torsional Strain in Crystal Structures of Small Molecules and Protein-Ligand Complexes using ab Initio Calculations. J Chem Inf Model 2019, 59, 4195–4208.

15. Perola, E.; Charifson, P. S., Conformational analysis of drug-like molecules bound to proteins: an extensive study of ligand reorganization upon binding. Journal of medicinal chemistry 2004, 47, 2499–2510.

16. Harder, E.; Damm, W.; Maple, J.; Wu, C.; Reboul, M.; Xiang, J. Y.; Wang, L.; Lupyan, D.; Dahlgren, M. K.; Knight, J. L., OPLS3: a force field providing broad coverage of drug-like small molecules and proteins. Journal of chemical theory computation 2016, 12, 281–296.

17. Kim, S.; Lee, J.; Jo, S.; Brooks III, C. L.; Lee, H. S.; Im, W., CHARMM‐GUI ligand reader and modeler for CHARMM force field generation of small molecules. Journal of computational chemistry 2017, 38, 1879–1886.

18. Wang, J.; Wolf, R. M.; Caldwell, J. W.; Kollman, P. A.; Case, D. A., Development and testing of a general amber force field. Journal of computational chemistry 2004, 25, 1157–1174.

19. Greenidge, P. A.; Kramer, C.; Mozziconacci, J. C.; Sherman, W., Improving docking results via reranking of ensembles of ligand poses in multiple X-ray protein conformations with MM-GBSA. J Chem Inf Model 2014, 54, 2697–717.

20. Fu, Z.; Li, X.; Merz Jr, K. M., Accurate assessment of the strain energy in a protein‐bound drug using QM/MM X ‐ ray refinement and converged quantum chemistry. Journal of computational chemistry 2011, 32, 2587–2597.

21. Schaerfer, C.; Schulz-Gasch, T.; Ehrlich, H. C.; Guba, W.; Rarey, M.; Stahl, M., Torsion Angle Preferences in Druglike Chemical Space: A Comprehensive Guide. Journal of Medicinal Chemistry 2013, 56, 2016–2028.

22. Guba, W.; Meyder, A.; Rarey, M.; Hert, J., Torsion Library Reloaded: A New Version of Expert-Derived SMARTS Rules for Assessing Conformations of Small Molecules. Journal of Chemical Information and Modeling 2016, 56, 1–5.

23. Groom, C. R.; Bruno, I. J.; Lightfoot, M. P.; Ward, S. C., The Cambridge structural database. Acta Crystallographica Section B: Structural Science, Crystal Engineering Materials 2016, 72, 171–179.

24. Berman, H. M.; Westbrook, J.; Feng, Z.; Gilliland, G.; Bhat, T. N.; Weissig, H.; Shindyalov, I. N.; Bourne, P. E., The protein data bank. Nucleic acids research 2000, 28, 235–242.

25. James, C.; Weininger, D.; Delany, J., In; Daylight Chemical Information Systems: Laguna Niguel, CA: 2000.

26. Schärfer, C.; Schulz-Gasch, T.; Rarey, M., TorsionAnalyzer: exploring conformational space. Journal of Cheminformatics 2013, 5.

27. Kellogg, E. H.; Leaver‐Fay, A.; Baker, D., Role of conformational sampling in computing mutation‐induced changes in protein structure and stability. Proteins: Structure, Function, Bioinformatics 2011, 79, 830–838.

28. Dunbrack Jr, R. L.; Cohen, F. E., Bayesian statistical analysis of protein side‐chain rotamer preferences. Protein Science 1997, 6, 1661–1681.

29. Coleman, R. G.; Carchia, M.; Sterling, T.; Irwin, J. J.; Shoichet, B. K., Ligand pose and orientational sampling in molecular docking. PloS one 2013, 8, e75992.

30. Landrum, G., Rdkit documentation. 2013, 1, 1–79.

31. Meng, E. C.; Shoichet, B. K.; Kuntz, I. D., Automated docking with grid‐based energy evaluation. Journal of computational chemistry 1992, 13, 505–524.

32. Sharp, K. A.; Friedman, R. A.; Misra, V.; Hecht, J.; Honig, B., Salt effects on polyelectrolyte–ligand binding: Comparison of Poisson–Boltzmann, and limiting law/counterion binding models. Biopolymers: Original Research on Biomolecules 1995, 36, 245–262.

33. Gallagher, K.; Sharp, K., Electrostatic contributions to heat capacity changes of DNA-ligand binding. Biophysical journal 1998, 75, 769–776.

34. Mysinger, M. M.; Shoichet, B. K., Rapid context-dependent ligand desolvation in molecular docking. Journal of chemical information modeling 2010, 50, 1561–1573.

35. Mysinger, M. M.; Carchia, M.; Irwin, J. J.; Shoichet, B. K., Directory of useful decoys, enhanced (DUD-E): better ligands and decoys for better benchmarking. Journal of medicinal chemistry 2012, 55, 6582–6594.

36. Lagarde, N.; Zagury, J. F.; Montes, M., Benchmarking Data Sets for the Evaluation of Virtual Ligand Screening Methods: Review and Perspectives. J Chem Inf Model 2015, 55, 1297–307.

37. Fan, H.; Schneidman-Duhovny, D.; Irwin, J. J.; Dong, G.; Shoichet, B. K.; Sali, A., Statistical potential for modeling and ranking of protein–ligand interactions. Journal of chemical information modeling 2011, 51, 3078–3092.

38. Lorber, D. M.; Shoichet, B. K., Hierarchical docking of databases of multiple ligand conformations. Curr Top Med Chem 2005, 5, 739–49.

