## Supplementary material for "Ligand Strain Energy in Large Library Docking": SI figures

**ASSOCIATED CONTENT**

†contributed equally

*corresponding author

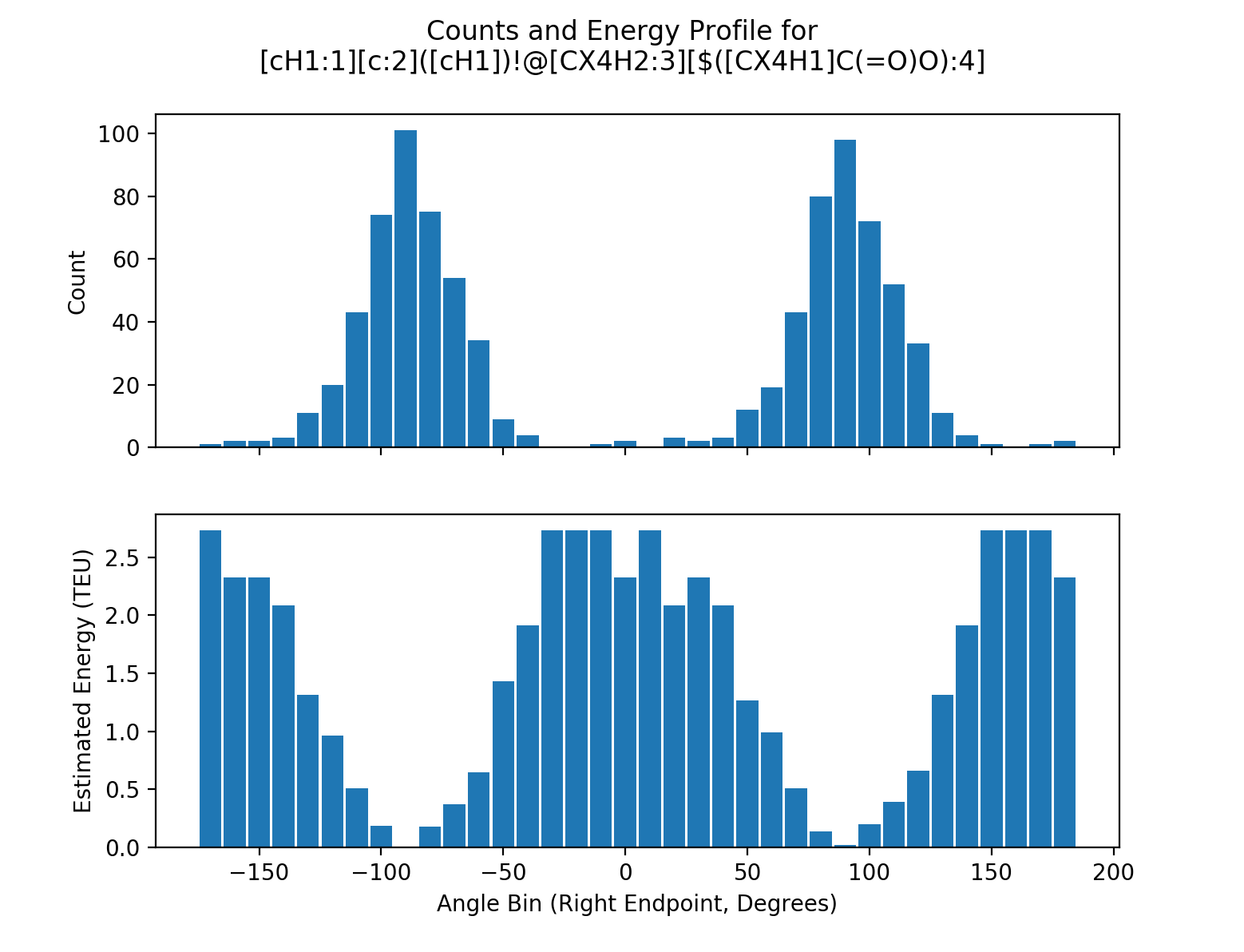

**Figure S1.** An example of torsion energy profile of a SMARTS pattern. The counts of each 10-degree bins are converted into torsion energy units (TEU).

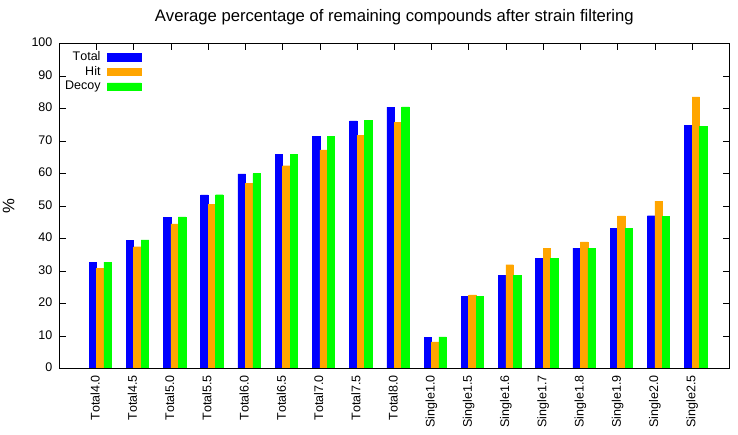

**Figure S2.** Average percentage of remaining compounds after strain filtering with different cut-offs. Blue is the total compounds in percentage, orange is the hits, and green is the decoys. Thresholds given in TEU.

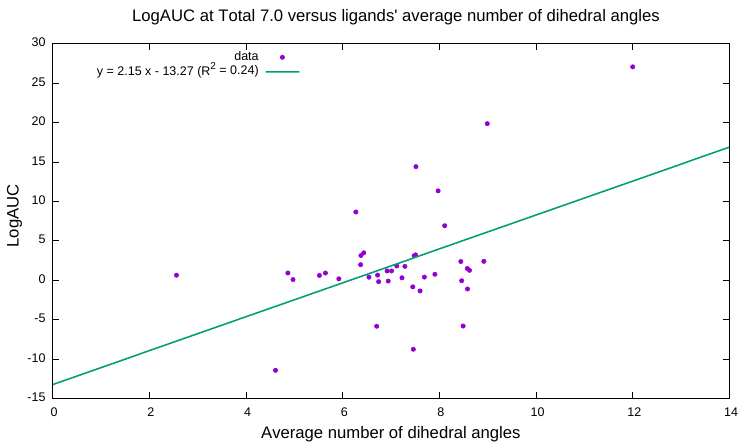

**Figure S3.** Correlation between ΔLogAUC and the ligands’ average number of dihedrals at the thresholds total strain 7.0 TEU. Each point represents a system in the DUD-E benchmark. There is a positive correlation between ΔLogAUC and the number of ligand and decoy rotatable bonds.

**Table S1.** Ligands docking score and strain energy in D4 receptor (Excel file)

**Table S2.** Ligands docking score and strain energy in AmpC (Excel file)

**Table S3.** DUD-E LogAUC results before and after strain filtering at different thresholds (Excel file)
